# Therapeutic Effect of Anti-FABP4 mAb (6H2) on Systemic Inflammation in Sepsis: Insights from Abdominal Organ Analysis

**DOI:** 10.1101/2025.04.16.649067

**Authors:** Muhammad Mustapha Ibrahim, Chunyan Li, Ma Yinzhong, Cheng Fang

## Abstract

Sepsis is a life-threatening condition driven by dysregulated immune responses and multi-organ dysfunction, with limited treatments targeting its underlying pathophysiology. Growing evidence highlights fatty acid-binding protein 4 (FABP4) as a key mediator of inflammation and organ injury in sepsis. In this study, we investigated the therapeutic potential of an anti-FABP4 monoclonal antibody (6H2) in a murine endotoxemia model induced by lipopolysaccharide (LPS). Treatment with 6H2 significantly attenuated systemic inflammation, as evidenced by modulated leukocyte responses, and provided substantial tissue protection in the liver, lungs, kidneys, and heart, reducing histopathological damage. Our findings identify 6H2 as a promising novel therapeutic intervention for sepsis.

**Author Summary:** **Title***Therapeutic Potential of Anti-FABP4 Antibody (6H2) in Sepsis: Protection Against Systemic Inflammation and Organ Damage*

*Key Findings:*

- Sepsis, a life-threatening inflammatory condition, lacks targeted therapies. We investigated **6H2**, a monoclonal antibody against fatty acid-binding protein 4 (FABP4), in a mouse model of sepsis.
- **6H2 treatment** significantly reduced sepsis severity, improved survival, and attenuated organ damage in the liver, lungs, kidneys, and heart.
- Histopathological analysis revealed **restored tissue integrity** in 6H2-treated mice, with reduced inflammation and cell death compared to controls.
- Complete Blood Count showed **modulated immune responses**, including altered leukocyte counts, suggesting 6H2’s role in rebalancing inflammation.

**Significance:** FABP4 is a key driver of sepsis-related inflammation and organ dysfunction. Our study demonstrates that **6H2** not only mitigates systemic inflammation but also provides **multi-organ protection**, offering a promising therapeutic strategy for sepsis. These findings support further preclinical and clinical development of FABP4-targeted therapies.

## Introduction

Sepsis remains a life-threatening condition characterized by a dysregulated host response to infection, leading to systemic inflammation, multi-organ dysfunction, and high mortality rates(1). Despite advances in critical care management, effective therapeutic strategies targeting the underlying inflammatory mechanisms are still urgently needed. Fatty acid-binding protein 4 (FABP4), also known as adipocyte FABP (A-FABP), has emerged as a key mediator of inflammation and metabolic dysfunction in various pathological conditions, including sepsis(2,3). Elevated FABP4 levels have been associated with increased cytokine production, endothelial dysfunction, and organ injury, suggesting its potential as a therapeutic target in sepsis (4).

Recent preclinical studies have demonstrated that monoclonal antibodies (mAbs) against FABP4, can attenuate systemic inflammation and improve outcomes in sepsis(5). However, the precise mechanisms by which anti-FABP4 mAb modulates inflammation in different abdominal organs— such as the liver, kidneys, and Lungs—remain incompletely understood. Given the critical role of these organs in sepsis-induced metabolic and immune dysregulation, a detailed analysis of their responses to FABP4 inhibition is essential for developing targeted therapies(6,7).

This study investigates the therapeutic potential of anti-FABP4 mAb (6H2) in a murine model of sepsis, with a focus on its effects on systemic inflammation and organ-specific responses. By evaluating cytokine profiles and histopathological changes, we provide novel insights into how Sepsis-Induced courses multiple organs damage. Our findings support the growing body of evidence that targeting FABPs could represent a promising strategy for reducing inflammation and improving survival in sepsis.

## Results

### 6H2 Treatment Mitigates Sepsis-Induced Physiological Dysfunction and Behavioral Alterations in a Murine Model

In the initial 24 hours following LPS injection, the surviving mice experienced considerable declines in body weight and deteriorated motor abilities (**Fig.1 B, C and D**). The observed deficits included decreased voluntary activity, erratic motion, irregular walking patterns, bristling of fur, partially shut eyelids, and slight eye discharge. These manifestations point towards systemic inflammation. We assessed the degree of sepsis using murine sepsis scores. The Sham group registered a low sepsis score at 2.5 ± 0.5, signifying the absence of significant systemic inflammation. Conversely, the LPS+mIgG group recorded a high sepsis score of 17.5 ± 1.2, verifying the successful induction of severe sepsis. However, the LPS+6H2 group had a substantially reduced sepsis score of 14.0 ± 1.5, indicating that 6H2 treatment could mitigate systemic inflammation to some extent. These results underscore that 6H2 can effectively reduce Murine Sepsis Score and mortality, modulate LPS-induced immune dysregulation, as evidenced by lower sepsis scores, and dampen systemic responses, thereby highlighting its therapeutic potential in inflammatory diseases.

**Figure 1:**
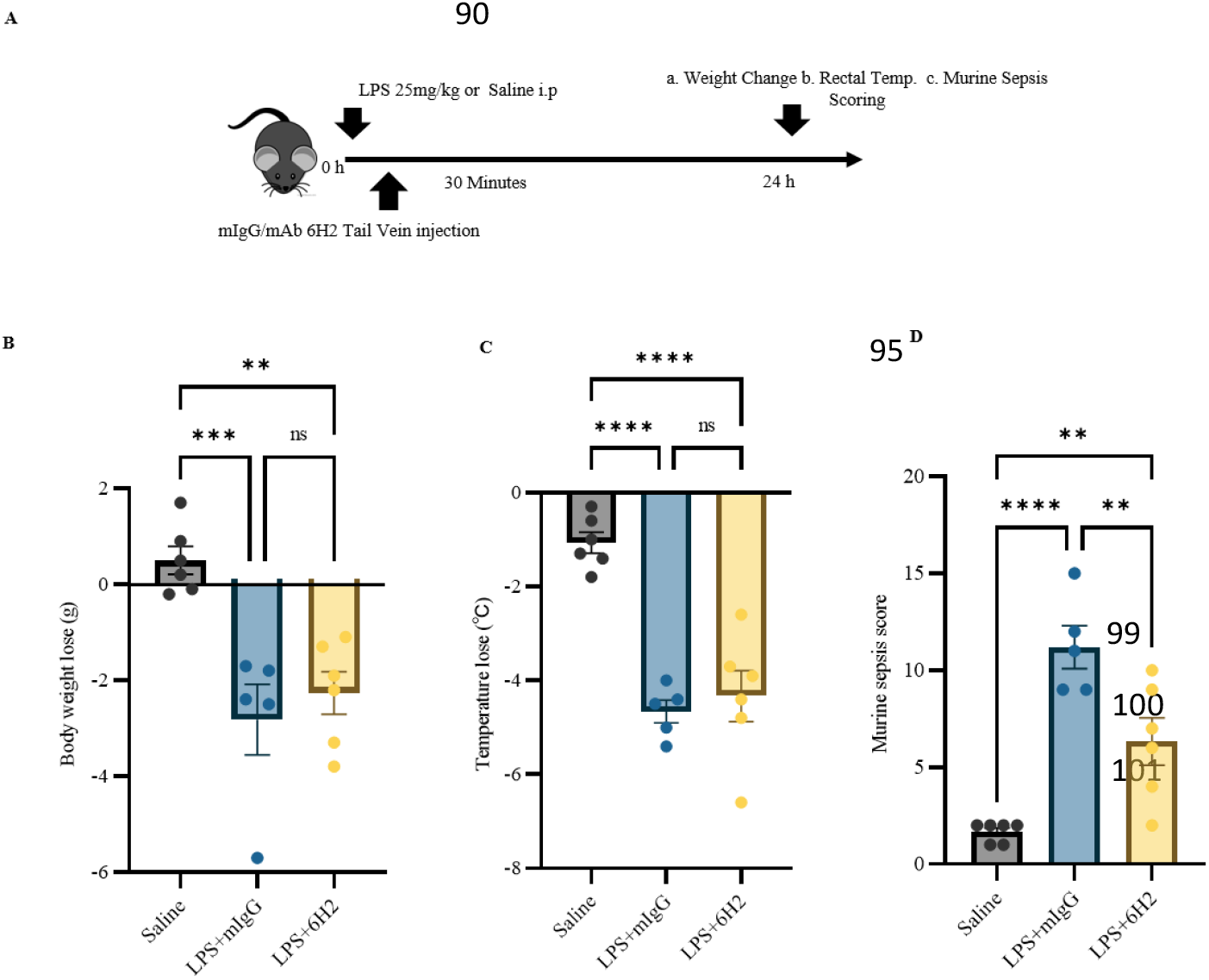
6H2 Treatment Reduces Mortality, Modulates Immune Responses, and Alleviates Systemic Effects in LPS-Induced Endotoxemia. (A) Experimental design: Endotoxemia was induced in male C57BL/6J mice (8-10 weeks) via LPS injection at the dose of (25 mg/kg body weight). (B) Body weight change at 24 h post-injection in saline (n = 5), LPS (n = 7), and 6H2 (n = 7) groups. (C) Loss of Rectal Temperature across the groups. (D) Murine sepsis scores reveal significant differences among Sham (2.5 ± 0.5), IgG (17.5 ± 1.2), and 6H2 (14.0 ± 1.5) groups.

### Therapeutic Administration of 6H2 Attenuates Systemic Inflammatory Responses in LPS-Induced Endotoxemia

In this study, we evaluated the therapeutic efficacy of 6H2 monoclonal antibody (mAb) in a murine model of LPS-induced endotoxemia using adult male C57BL/6J wild-type mice. The experimental design comprised three groups: saline-treated controls, LPS-challenged mice receiving isotype control antibody (mIgG, 3.6 mg/kg), and LPS-challenged mice treated with 6H2 mAb (3.6 mg/kg). The Complete Blood Count (CBC) analysis revealed a significant reduction in white blood cells (WBCs), including monocytes, neutrophils, and lymphocytes, within the LPS+mIgG group (**Fig. 2A**). This pattern indicates the presence of both acute and chronic inflammatory responses. Conversely, treatment with 6H2 substantially decreased overall leukocyte (WBC) counts while selectively altering the composition of leukocyte subsets. Specifically, 6H2 treatment led to an increase in monocytes and neutrophils while simultaneously reducing lymphocyte levels. This unique shift in immune cell populations suggests a potent immunomodulatory effect, which may promote the resolution of acute inflammation through the recruitment of myeloid cells and suppression of lymphoid activity.

**Fig. 2:**
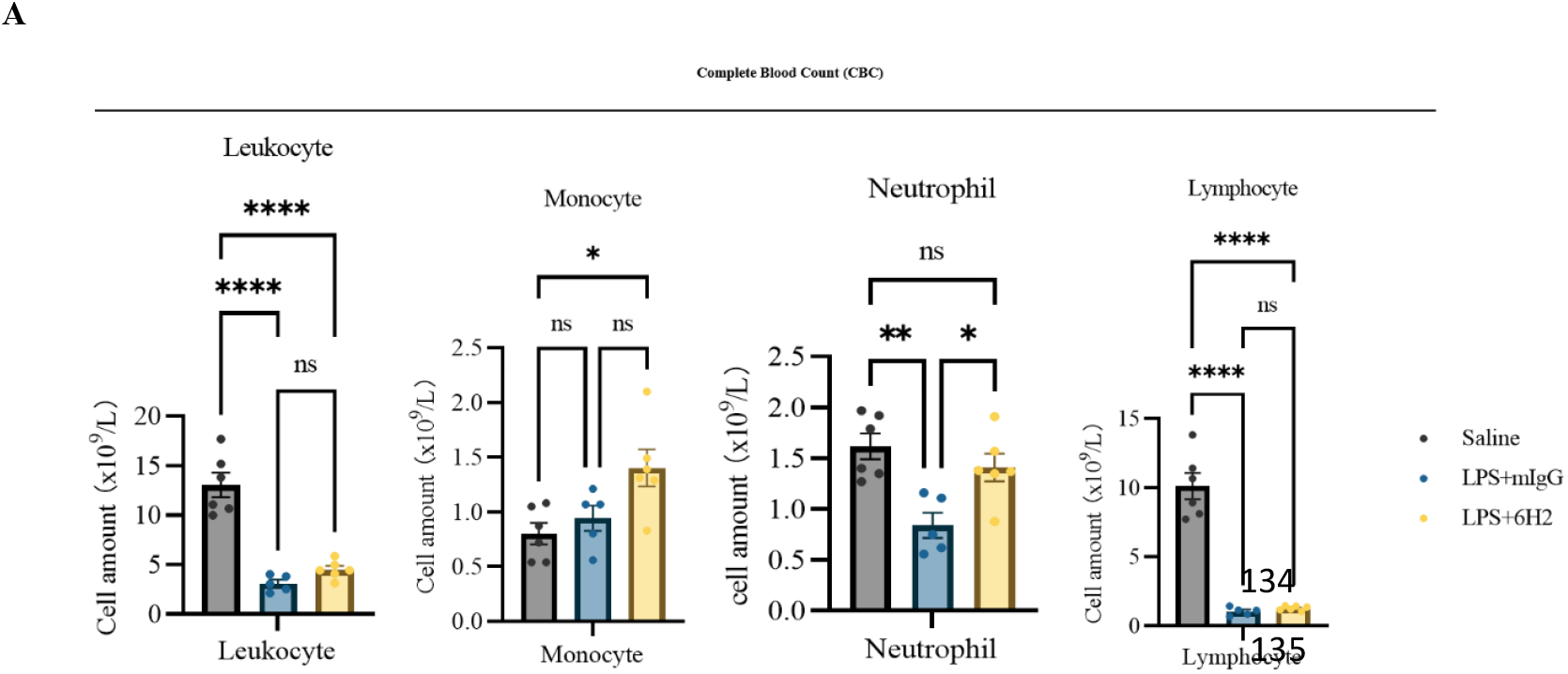
Systemic Responses to 6H2 Therapy in LPS-Induced Endotoxemia. (A) complete blood count profile

These findings indicate that 6H2 can effectively modulate LPS-induced immune dysregulation and mitigate systemic inflammatory responses, thereby highlighting its therapeutic potential for treating inflammatory diseases.

### Efficacy of 6H2 on Histopathological Assessment of Multiple Organs and Fabp4 Levels in LPS-Induced Endotoxemia

In this study, we evaluated the efficacy of 6H2 treatment on histopathological changes in multiple organs and systemic inflammation in a murine model of LPS-induced endotoxemia. Histopathological analysis (H&E staining) of organs, including the liver, lungs, kidneys, spleen, and heart, revealed significant tissue damage and inflammatory infiltration in LPS-treated mice compared to saline controls. However, 6H2 treatment markedly reduced these pathological changes, demonstrating its protective effects against LPS-induced organ damage (**Fig. 3A**).

**Fig. 3.**
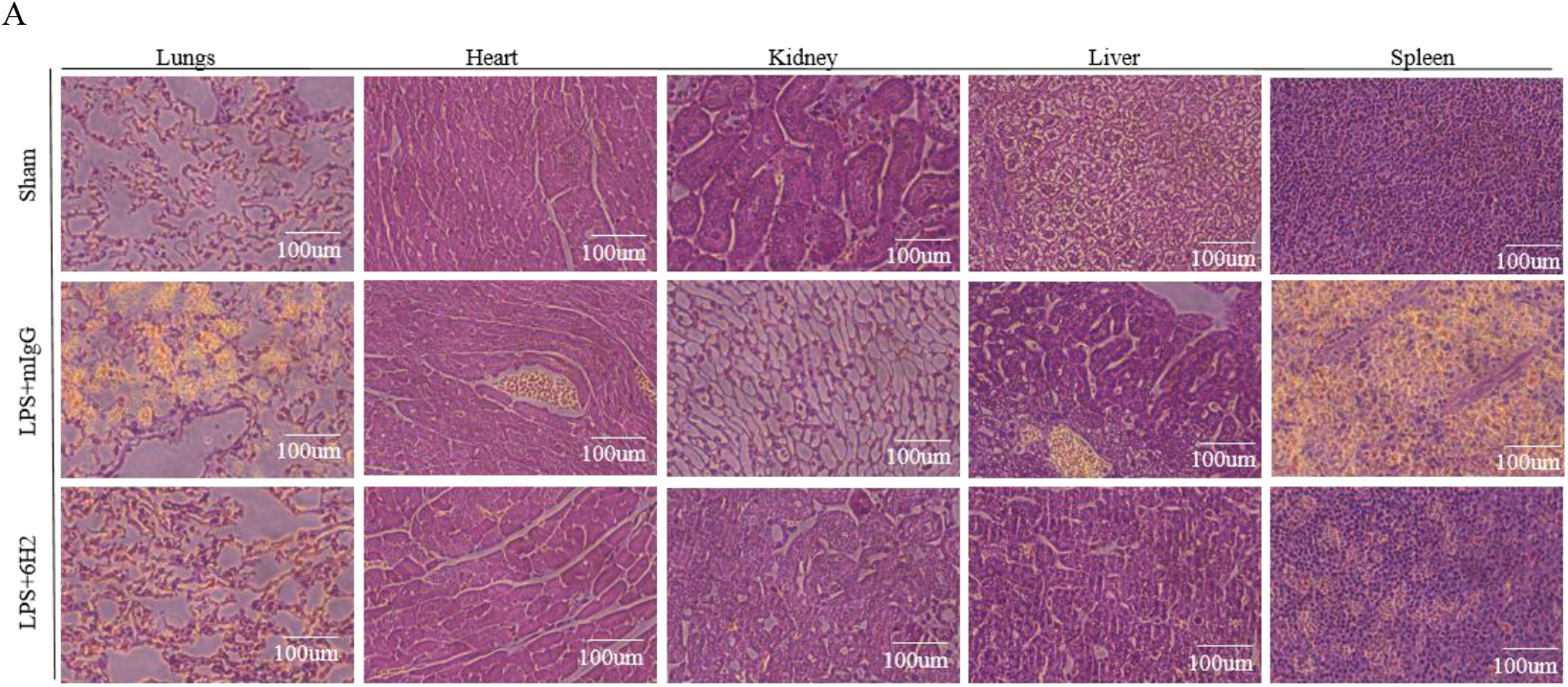
Impact of 6H2 on Tissue Morphology (H&E) and Multiple Organs Post Saline/LPS-Induced Inflammation. (A) Histopathological analysis reveals that LPS causes substantial tissue damage in various organs and 6H2 attenuates.

In the histopathological assessment of lung tissue, the sham group exhibited thin-walled alveoli with capillaries, air-filled alveolar spaces, and minimal interstitium. In contrast, the LPS+mIgG group showed thickened alveolar walls and altered alveolar spaces, indicative of damage. However, the LPS+6H2 group demonstrated a recovery process in both the epithelial and endothelial cells, with reduced thickening of alveolar walls and more normal alveolar spaces.

For liver tissue, the sham group displayed a typical lobular architecture with hepatocytes radiating from the central vein, eosinophilic cytoplasm, and sinusoids lined by endothelial cells. The LPS+mIgG group exhibited hepatocyte swelling, fatty changes, inflammation, and cirrhosis. In contrast, the LPS+6H2 group showed restored architectural lobular and hepatocyte structures with clear cytoplasm, indicating a recovery process.

In kidney tissue, the sham group had glomeruli with dark blue nuclei and a central Bowman’s capsule, with tubules lined by cuboidal cells. The LPS+mIgG group displayed glomerulosclerosis, tubular degeneration, interstitial inflammation, and acute injury. The LPS+6H2 group showed a healing process in the central Bowman’s capsule and kidney cortex, with improved glomerular and tubular structures.

Heart tissue in the sham group was well organized with clear cross-striations and centrally located nuclei. The LPS+mIgG group exhibited disorganized muscle fibers, eosinophilic necrosis, and infiltration of inflammatory cells. The LPS+6H2 group showed restored well-organized myocardial fibers with clear striations, indicating a recovery process.

The spleen in the sham group had white pulp with germinal centers containing proliferating lymphocytes and red pulp with sinusoids. The LPS+mIgG group showed congestion and infarction. The LPS+mAb6H2 group exhibited restoration and proliferating lymphocytes, indicating a recovery process.

Together, these findings highlight the potential of 6H2 in mitigating LPS-induced organ damage and systemic inflammation. The histopathological improvements observed across multiple organs indicate that 6H2 has a broad-spectrum protective effect, making it a promising therapeutic candidate for treating LPS-induced endotoxemia.

## Discussion

Our findings that 6H2 treatment can mitigate the physiological and behavioral alterations induced by LPS in a murine model are supported by several lines of scientific evidence(8). The observed symptoms, such as reduced spontaneous movement, erratic motion, and piloerection, are consistent with the hallmarks of systemic inflammation and sepsis in murine models. The use of murine sepsis scores to evaluate the severity of sepsis is a well-established method, with studies showing that these scores can effectively differentiate between mild and severe sepsis(9,10). The significant reduction in sepsis scores in the LPS+6H2 group compared to the LPS+mIgG group indicates that 6H2 treatment can modulate the inflammatory response and alleviate the severity of sepsis(11).The ability of 6H2 to reduce mortality and modulate immune dysregulation aligns with the broader understanding of sepsis pathophysiology. Sepsis is characterized by a dysregulated immune response that leads to widespread inflammation and organ dysfunction(12). The observed reduction in sepsis scores and systemic responses in the LPS+6H2 group suggests that 6H2 may act by modulating key inflammatory pathways. This is supported by studies showing that interventions targeting specific inflammatory mediators can improve outcomes in sepsis models

The modulation of leukocyte populations by 6H2—specifically, the increase in monocytes and neutrophils alongside a reduction in lymphocytes—points to a unique immunomodulatory mechanism. This shift may reflect enhanced myeloid cell recruitment to sites of inflammation, promoting tissue repair while suppressing excessive lymphoid activation. Such a response could be beneficial in sepsis, where immune dysregulation often leads to simultaneous hyperinflammation and immunosuppression. These findings are consistent with recent work showing that FABP4 influences macrophage polarization and cytokine release, further underscoring its role in immune cell regulation(13,14).

Histopathological analysis revealed that 6H2 treatment significantly mitigated organ damage in the liver, lungs, kidneys, and heart. In the liver, 6H2 restored lobular architecture and reduced hepatocyte swelling, which contrasts sharply with the cirrhosis and fatty changes seen in LPS-treated controls. Similar protective effects were observed in the lungs, where 6H2 attenuated alveolar wall thickening and preserved airspace integrity. These improvements correlate with prior studies demonstrating that FABP4 inhibition reduces oxidative stress and tissue injury in metabolic and inflammatory diseases(15). The kidney findings are particularly noteworthy, as 6H2 ameliorated glomerulosclerosis and tubular degeneration, suggesting a potential role for FABP4 in sepsis-associated acute kidney injury.

In conclusion, our data strongly support the therapeutic potential of 6H2 in sepsis by demonstrating its ability to reduce systemic inflammation and protect against multi-organ damage. These findings build upon emerging evidence that FABP4 is a viable target for inflammatory diseases and provide a foundation for future preclinical and clinical studies.

## Materials and Methods

Male C57BL/6J mice (8–10 weeks old) were obtained from Vital River Laboratory (Beijing, China) and maintained under pathogen-free conditions with a 12-hour light/dark cycle and free access to food and water, in accordance with NIH guidelines (NIH Publication No. 86-23, revised 1985). All experimental procedures were approved by the Institutional Animal Care and Use Committee (IACUC) of the Shenzhen Institute of Advanced Technology, Chinese Academy of Sciences. To induce endotoxemia, mice received an intraperitoneal (i.p.) injection of lipopolysaccharide (LPS; Escherichia coli O55:B5, Sigma-Aldrich, #L2880) at a dose of 25 mg/kg. Thirty minutes after LPS administration, animals were randomly assigned to receive an intravenous injection of either anti-FABP4 monoclonal antibody (6H2; 3.6 mg/kg) or an isotype control antibody (mIgG; Immuno-Diagnostics, #221116, RRID: AB_3073816) at an equivalent dose. Mice that succumbed to sepsis prior to the predetermined experimental endpoint were excluded from subsequent analyses.

## Behavioral test

### Development of the MSS

In a preliminary experiment, 30 mice were allocated into three groups (Sham control, IgG, and 6H2), with sepsis induced in the IgG and 6H2 groups via LPS administration (25 mg/kg). After 24 hours, rectal temperature and body weight measurements revealed hypothermia (pharexia) and an initial weight decline followed by slight recovery. Sepsis severity was assessed using the Murine Sepsis Score (MSS), which evaluated spontaneous activity, response to stimuli, posture, respiratory rate, breathing quality, and piloerection (each scored 0–4; **Table 1**), providing a standardized measure of sepsis progression and treatment effects.

**Table 1.** Murine Sepsis Score (MSS) to assess the severity of disease in an experimental model of sepsis induced by LPS.

| *Variable* | *Score and description* |
| --- | --- |
| **Appearance** | 0. Coat is smooth<br>1. Patches of hair piloerected<br>2. The majority of the back is piloerected<br>3. Piloerection may or may not be present, the mouse appears “puffy”<br>4. Piloerection may or may not be present, mouse appears emaciated |
| **Level of consciousness** | 0. Mouse is active<br>1. The mouse is active but avoids standing upright<br>2. Mouse activity is noticeably slowed. The mouse is still ambulant.<br>3. Activity is impaired. Mouse only moves when provoked, movements have a tremor<br>4. Activity severely impaired. Mouse remains stationary when provoked, with possible tremor |
| **Activity** | 0. Normal amount of activity. Mouse is any of: eating, drinking, climbing, running, fighting<br>1. Slightly suppressed activity. The mouse is moving around the bottom of the cage<br>2. Suppressed activity. The mouse is stationary with occasional investigative movements<br>3. No activity. Mouse is stationary<br>4. No activity. Mouse experiencing tremors, particularly in the hind legs |
| **Response to stimulus** | 0. The Mouse responds immediately to auditory stimulus or touch<br>1. Slow or no response to auditory stimulus; strong response to touch (moves to escape)<br>2. No response to auditory stimulus; moderate response to touch (moves a few steps)<br>3. No response to auditory stimulus; mild response to touch (no locomotion)<br>4. No response to auditory stimulus. Little or no response to touch. It cannot right itself if pushed over |
| **Eyes** | 0. Open<br>1. Eyes not fully open, possibly with secretions<br>2. Eyes at least half closed, possibly with secretions<br>3. Eyes half closed or more, possibly with secretions<br>4. Eyes closed or milky |
| **Respiration rate** | 0. Normal, rapid mouse respiration<br>1. Slightly decreased respiration (rate not quantifiable by eye)<br>2. Moderately reduced respiration (rate at the upper range of quantifying by eye)<br>3. Severely reduced respiration (rate easily countable by eye, 0.5 s between breaths)<br>4. Extremely reduced respiration (>1 s between breaths) |
| **Respiration quality** | 0. Normal<br>1. Brief periods of labored breathing<br>2. Laboured, no gasping<br>3. Laboured with intermittent gasps<br>4. Gasping |

## Histological Analysis

Tissue samples were fixed in 10% neutral-buffered formalin (NBF) for 24-36 hours to ensure optimal morphological preservation and prevent autolysis. Following fixation, tissues were rinsed and stored in 70% ethanol until processing. Organs including heart, liver, lungs, kidneys, and spleen were dissected, with dense tissues undergoing decalcification in ethylenediaminetetraacetic acid (EDTA) when necessary. Tissues were then dehydrated through a graded ethanol series (70%-100%), cleared in xylene, and embedded in paraffin wax. Serial 2-μm sections were cut using a precision microtome and mounted on glass slides. For morphological assessment, sections were stained with hematoxylin and eosin (H&E): hematoxylin (nuclear stain; blue) highlighted cellular nuclei and basophilic structures, while eosin (cytoplasmic stain; pink) delineated cytoplasmic and extracellular matrix components, enabling comprehensive evaluation of tissue architecture and pathological changes.

## Complete Blood Count (CBC) Analysis

Blood samples were collected via terminal cardiac puncture immediately prior to perfusion to ensure optimal sample quality and prevent hemodilution. Using aseptic technique, approximately 200 μL of whole blood was drawn from each mouse using a sterile 25-gauge needle and syringe pre-coated with EDTA anticoagulant. The blood was immediately transferred to EDTA-treated microtainer tubes (BD Biosciences) and gently inverted 8-10 times to ensure proper mixing with the anticoagulant and prevent clot formation.

For comprehensive hematological profiling, samples were analyzed within 2 hours of collection using an automated veterinary hematology analyzer (e.g., IDEXX ProCyte Dx). The CBC panel included quantification of Erythrocyte, Leukocyte differential count (neutrophils, lymphocytes, monocytes). Prior to analysis, instrument calibration was verified using manufacturer-provided control samples. Samples showing signs of hemolysis or microclots were excluded from analysis. This standardized protocol ensured reliable assessment of systemic inflammatory responses and hematological alterations associated with sepsis.

## Statistical Analysis

Quantitative data are expressed as mean ± standard error of the mean (SEM) to represent both central tendency and measurement precision. All statistical analyses were conducted using GraphPad Prism software (version 10.1.0, GraphPad Software Inc.). For comparisons between two experimental groups, we employed a two-tailed unpaired Student’s t-test with Welch’s correction to account for potential variance inequality. Multi-group comparisons were performed using one-way analysis of variance (ANOVA) followed by Tukey’s post hoc test for multiple comparisons when statistical significance was detected.

The threshold for statistical significance was established at *P* < 0.05. For all ANOVA analyses, we first confirmed the assumptions of normality (using Shapiro-Wilk test) and homogeneity of variance (using Brown-Forsythe test). In cases where these assumptions were violated, appropriate non-parametric alternatives (Mann-Whitney U test for two groups or Kruskal-Wallis test with Dunn’s post hoc for multiple groups) were substituted.

## Acknowledgements

The authors are grateful

## Author contributions

Muhammad Mustapha Ibrahim and Chunyan Li performed the research. Muhammad Mustapha Ibrahim and Cheng Fang analyzed the experimental data and wrote the manuscript. Cheng Fang. Supervised the laboratory experiments. All other authors Reviewed and commented the manuscript.

## Funding

The funders were not involved in study design, data collection, data analysis, Manuscript preparation and/or publication decisions.

## Data availability

No datasets were generated or analyzed during the current study.

## Declarations

## Ethics approval and consent to participate

All animal work was approved by the Institutional Animal Care and Use Committee of Shenzhen Institute of Advanced Technology, Chinese Academy of Sciences.

## Consent for publication

All authors agreed on the content of this manuscript before its submission.

## Conflict interests

The authors declare no conflicts interests.

## Notes

### Competing Interest Statement

The authors have declared no competing interest.

